# Phenotypic profile of *Mycobacterium tuberculosis*-specific CD4 T cell responses in HIV-positive patients who develop Tuberculosis-associated Immune Reconstitution Inflammatory Syndrome

**DOI:** 10.1101/2022.07.20.500909

**Authors:** Raymond M Moseki, Daniel L Barber, Elsa Du Bruyn, Muki Shey, Helen Van der Plas, Robert J Wilkinson, Graeme Meintjes, Catherine Riou

## Abstract

**Background:** Tuberculosis-associated immune reconstitution inflammatory syndrome (TB-IRIS) is a frequent complication of co-treatment for TB and HIV-1. We characterized Mtb-specific CD4 T cell phenotype and transcription factor profile associated with the development of TB-IRIS.

**Methods:** We examined the role of CD4 T-cell transcription factors in a murine model of mycobacterial IRIS. In humans, we compared longitudinally on antiretroviral therapy (ART) the magnitude, activation, transcription factor profile and cytotoxic potential of Mtb-specific CD4 T cells between TB-IRIS (n=25) and appropriate non-IRIS control patients (n=18) using flow cytometry.

**Results:** In the murine model, CD4 T cell expression of Eomes, but not Tbet, was associated with experimentally induced IRIS. In patients, TB-IRIS onset was associated with the expansion of Mtb-specific IFNγ+CD4 T cells (p=0.039). TB-IRIS patients had higher HLA-DR expression (p=0.016), but no differences in the expression of T-bet or Eomes were observed. At TB-IRIS onset, Eomes+Tbet+Mtb-specific IFNγ+CD4+ T cells showed higher expression of Granzyme B in TB-IRIS patients (p=0.026).

**Conclusion:** While the murine model of MAC-IRIS suggests that Eomes+CD4 T cells underly IRIS, TB-IRIS was not associated with Eomes expression in patients. Mtb-specific IFNγ+CD4 T cell responses in TB-IRIS patients are differentiated, highly activated and potentially cytotoxic.

## BACKGROUND

Although antiretroviral therapy (ART) has substantially reduced HIV-1 related morbidity and mortality in patients with HIV-associated tuberculosis (TB) [1], TB-immune reconstitution inflammatory syndrome (TB-IRIS) frequently complicates management [2, 3]. TB-IRIS has an estimated incidence of 18% across cohorts and an attributable mortality rate of 2% [4].

Two forms of TB-IRIS are recognised: 1) Unmasking TB IRIS which occurs in patients with undiagnosed TB who present with severe inflammatory features of TB during the first 3 months of ART and 2) Paradoxical TB-IRIS which occurs in patients started on TB treatment before ART who experience recurrent, new or worsening symptoms and signs of TB within the first months of initiating ART [5, 6]. The major risk factors for paradoxical TB-IRIS include a low CD4 count prior to ART initiation, higher HIV-1 viral load, a short interval between TB treatment and ART initiation and disseminated TB [7, 8].

Innate immune responses including inflammasome activation [9, 10], monocyte and natural killer cell activation [11, 12], neutrophilia [12, 13] and dysregulation of the complement system in monocytes [14] have been associated with TB-IRIS. Elevated concentrations of proinflammatory cytokines [15, 16] and matrix degrading metalloproteinases [17] have been described at TB-IRIS onset. Moreover, monocyte subset frequency and circulating inflammatory mediators can independently predict TB-IRIS disease [18, 19]. Expansion of pathogen-specific CD4+ T cells has been observed in association with TB-IRIS [20–23]. Pathogen-specific CD4+ T cells from patients with IRIS have been reported to be highly activated [24] and polyfunctional [25]. Recently, it was reported that HIV-1 patients with *Mycobacterium avium* complex (MAC) infection, who developed MAC-IRIS had higher expression of Eomesodermin (Eomes) compared with Tbet in MAC-specific IFNγ+CD4+ T cells at the onset of IRIS [26]. Eomes and Tbet are members of the T-box DNA binding family of transcription factors with structural similarities and overlapping expression [27]. Eomes is involved in the development of cytotoxic T lymphocyte activity [27] while Tbet is a Th-1 lineage-defining transcription factor [28]. Th-1 responses have been implicated in a mouse model of MAC-IRIS [20, 21]. Consequently, we capitalized on the existing mouse model of IRIS to investigate phenotypic CD4 T cell features that may be associated with IRIS in mice and compare these with findings in patients developing TB-IRIS in a prospective cohort study of patients with HIV-associated TB initiating ART.

## METHODS

### *M. avium*-IRIS induction in mice

C57BL/6J-[KO] TCRalpha mice (6-8 weeks old) were intravenously infected with 1×10^6^ colony-forming units of *M. avium* (strain SmT 2151). After at least 40 days, CD4 T cells were isolated from C57BL/6, B6.129S6-Tbx21tm1Glm/J mice (The Jackson Laboratory, Bar Harbor, ME), or eomes^fl/fl^CD4-CRE^+^ uninfected mice using positively selecting microbeads (Miltenyi Biotec, Auburn, CA), and 1×10^6^ cells were intravenously transferred into chronically infected T cell deficient mice. All mice were maintained and bred at NIAID, NIH, Bethesda, MD. All animals were housed at an Association for the Assessment and Accreditation of Laboratory Animal Care-approved facility at the NIAID according to the National Research Council’s Guide for the Care and Use of Laboratory Animals

### Participants in clinical study

Samples were obtained from patients with HIV-associated TB initiating ART enrolled in a prospective observational study conducted at Brooklyn Chest Tuberculosis Hospital between May 2009 and November 2010 in Cape Town, South Africa [29]. All patients were ART naïve and those with rifampicin-resistant TB were excluded. TB diagnosis was based on smear, culture or clinical diagnosis. The first TB episode was treated with standard first line regimen of rifampicin (R), isoniazid (H), pyrazinamide and ethambutol for two months followed by four months of RH regimen. In patients with subsequent episodes, streptomycin was added for 2 months. TB-IRIS was diagnosed per International Network for the Study of HIV-associated IRIS (INSHI) criteria [5]. HIV-1 treatment included lamivudine and efavirenz with stavudine or tenofovir depending on guidelines at the time. Written informed consent was obtained from all participants. The study was approved by the Human Research Ethics Committee (HREC REF: 049/2009 and 809/2018) of the University of Cape Town. Clinical and other immunological findings from this cohort have been published [9, 29, 30].

### PBMC isolation and stimulation

PBMC were isolated by Ficoll-Hypaque density gradient centrifugation (GE Healthcare® ALC-PK121R), cryopreserved and stored. Cryopreserved PBMC were thawed and rested at 37 °C in RPMI 1640 containing 10% heat-inactivated FCS for 4 hours prior to antigen stimulation. PBMC (2×10^6^ cells) were stimulated with a peptide pool constituted of 300 Mtb-derived peptides (Mtb300, 1.5 µg/mL) [31] in the presence of anti-CD28 and anti-CD49d antibodies (both at 1 μg/ml, BD, Franklin Lakes, New Jersey) and brefeldin-A (10 μg/ml, Sigma, St Louis, Missouri) for 6 hours. Unstimulated cells, incubated with co-stimulatory antibodies and Brefeldin A only, were used as controls.

### Cell staining and acquisition

Samples with a cell count of less than one million or a viability score of less than 20% were excluded. After stimulation, cells were washed, stained with a viability marker (Live/Dead® Fixable Near-InfraRed marker, Invitrogen, Carlsbad, California) for 10 minutes at room temperature and subsequently surface stained with the following antibodies: CD4-PerCP-cy5.5, PD1-BV421, HLA-DR-BV605, CD14-Allophycocyanin/H7, CD19-Allophycocyanin/H7 (all from Biolegend, San Diego, California) and CD8-Alexa700 (BD, Franklin Lakes, New Jersey) for 30 minutes at room temperature. Cells were fixed and permeabilized using the eBioscience Foxp3 fixation buffer for 30 minutes at room temperature and stained with IFNγ-BV711 (Biolegend), TNFα-FITC (Biolegend), granzyme B-BV510 (BD), Eomes-eFluor 660 (e-Bioscience, San Diego, California), Tbet-PEcy7 (e-Bioscience) and CD3-BV785 (Biolegend), for 45 minutes at 4 °C. Cells were washed and resuspended in 1% formaldehyde in PBS. Samples were acquired in the BD LSRII and data were analysed using FlowJo software version 9.9.6 (BD). The gating strategy is presented in **Supplementary Figure 1**. A positive Mtb-specific IFNγ response was defined as three-fold higher than the background measured in the presence of co-stimulatory antibodies without antigen. For the phenotypic analyses of Mtb-specific IFNγ+CD4+ T cells, only Mtb responses with more than 20 events were considered. Protocols were compliant with the guidelines for flow cytometry in immunological studies [32]. Although we analysed immunological characteristics of live cells, our cohort included 8 samples with a viability below 50% (median: 67%, [range: 96-22%]). Prior to assessing immunological phenotypic characteristics of our cohort, we ascertained whether sample viability affected immunological expression of markers (particularly Tbet and Eomes) in our cohort. There was no correlation between sample viability and the expression of Tbet and Eomes at all measured time points (data not shown).

### Statistical analyses

For analyses, samples from IRIS and non-IRIS groups were classified into four categories based on sample timing in relation to ART: Baseline (BL) include samples collected within seven days before or on the day of ART initiation, Week 2 (W2)-samples collected between day 1 and 14 on ART, Week 4 (W4)-samples collected between day 15 and 30 on ART, Week 6 (W6) samples collected between day 31 and 65 on ART (Supplementary Table 2&3). Paired samples were analysed using the paired Wilcoxon ranked Student T test while the Mann-Whitney U test was used to compare unpaired samples for all time points between TB-IRIS and non-IRIS groups. A p-value of 0.05 or less was considered statistically significant. All statistical analyses were performed using Prism (v8.0.2, GraphPad Software Inc, San Diego, CA, USA).

## RESULTS

### Role of Eomes and Tbet in CD4 T cells during experimentally-induced IRIS

To model IRIS, T cell deficient (TCRα-/-) mice were infected with *M. avium*. This reproduced a lymphopenic host harbouring a mycobacterial infection. After 40-60 days, the mice were injected with CD4 T cells to mimic the reconstitution of T cells that occurs after ART **(Figure 1A)**. To examine the expression of Eomes and Tbet in CD4 T cells and their potential involvement in the murine model of IRIS, we injected mice with WT, Tbet^-/-^, and Eomes deficient CD4 T cells and examined the donor CD4 T cells ten days post injection. We found that during murine IRIS, CD4 T cells surprisingly expressed little Tbet. Instead, a significant population of Eomes+ CD4 T cells was observed **(Figure 1B)**. WT and Tbet^-/-^ CD4 T cells induced similar levels of weight loss **(Figure 1C)**. In contrast, recipients of Eomes deficient CD4 T cells displayed less weight loss and longer survival compared to mice injected with WT CD4 T cells **(Figure 1D, E)**. We concluded that, CD4 T cells utilize Eomes but not Tbet, to drive *M. avium* IRIS in this animal model. These findings prompted us to next examine the role of Eomes expressing CD4 T cells in human TB-IRIS.

**Figure 1.**
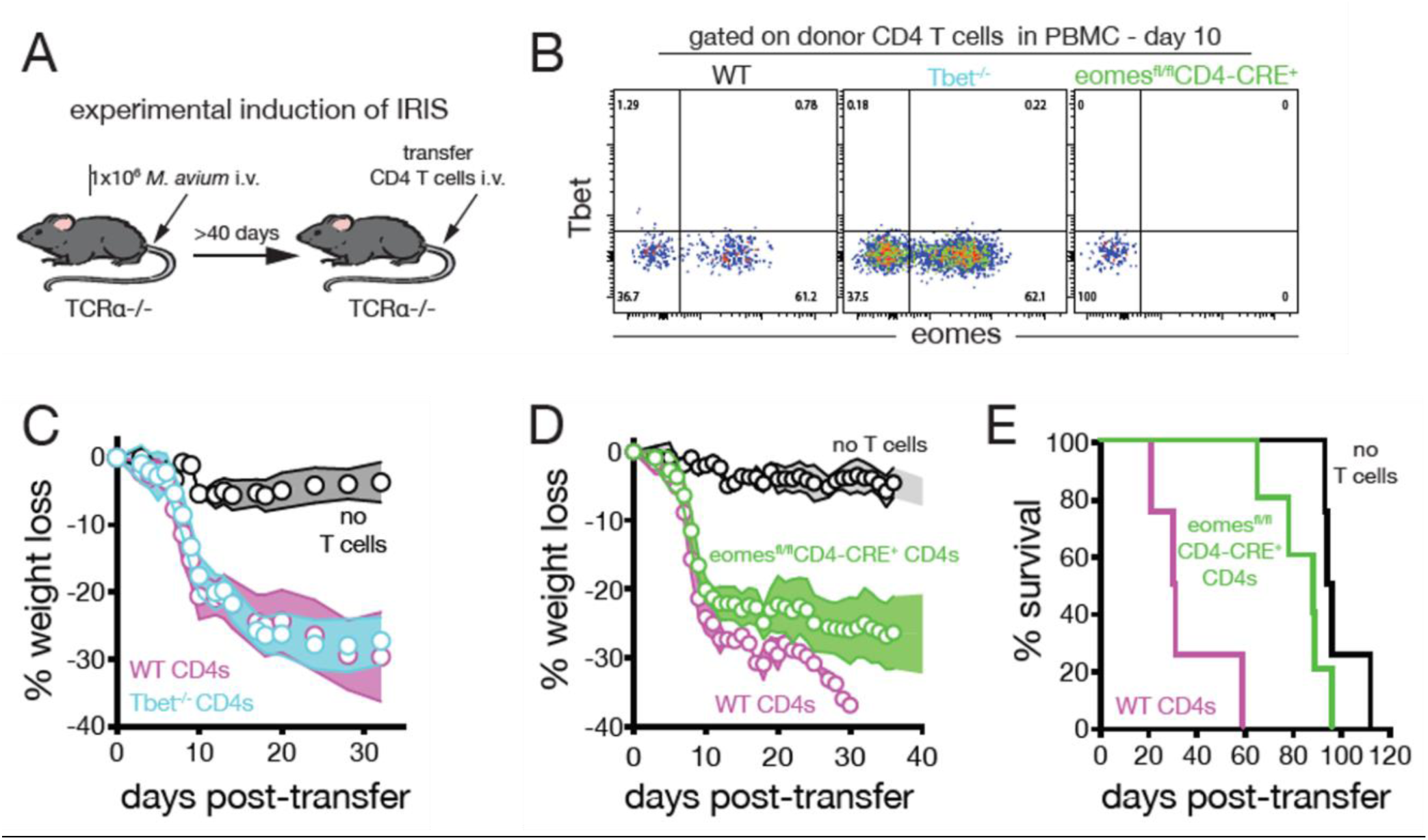
CD4 T cell expression of eomesodermin promotes mycobacterial IRIS in a murine model. **A**. To model IRIS in mice, TCRα-/- mice harboring a chronic *M. avium* infection were injected with purified CD4 T cells from uninfected donor mice. **B**. *M. avium* infected TCRα-/- mice were injected with WT, Tbet-deficient or eomes-deficient CD4 T cells. The donor CD4 T cells (CD4+TCRβ+CD3+) were analyzed in the PBMC on day 10 post infection for the expression of Tbet and eomesodermin (plots are concatenated from n=8 mice/group). **C**. *M. avium* infected TCRα-/- mice were injected with either WT or Tbet-deficient CD4 T cells and monitored for weight loss (n=5 mice/group). **D**. *M. avium* infected TCRα-/- mice were injected with no T cells, WT or eomes-deficient CD4 T cells and monitored for weight loss. **E**. Survival of mice receiving WT or eomes-deficient CD4 T cells. n=4 to 5 mice/group. Error bars represent the standard deviation. Data are representative of at least 4 independent experiments each.

### Clinical characteristics of the cohort

Sufficient samples for immunological analyses were available for 43 HIV-1 infected inpatients being treated for TB when starting ART: 25 patients developed TB-IRIS and 18 patients did not (non-IRIS controls). The demographic and clinical characteristics of the two groups are summarized in **Table 1**. In both groups, over three-quarters of patients had evidence of extrapulmonary TB and around 20% had neurological TB, a common reason for TB patients in South Africa to be admitted to a TB hospital. Notably, 7/25 (28%) of TB-IRIS and 4/18 (22%) of non-IRIS patients were on treatment with corticosteroids at the time of starting ART, the most frequent indication being neurological TB. We previously demonstrated that corticosteroid therapy had no significant effect on the frequency of Mtb-specific CD4 T cells [33]. The median CD4 count at the start of ART was lower in TB-IRIS patients (median: 68 cells/mm^3^) compared with non-IRIS patients (median 111 cells/mm^3^) (p=0.009). The median duration of TB treatment before initiation of ART was similar for the groups (median 37 days in TB-IRIS versus 32 days in non-IRIS patients). The duration of ART prior to developing TB-IRIS symptoms was a median of 15 days. Additional clinical data is presented in **Supplementary Table 1**.

**Table 1.** Clinical characteristics of patients who developed tuberculosis immune reconstitution inflammatory syndrome (TB-IRIS, n=25) and those who did not (non-IRIS, n=18). TB: Tuberculosis, PTB: pulmonary TB, EPTB: extrapulmonary TB, IQR: interquartile range, ART: antiretroviral treatment, Hb: hemoglobin. The Wilcoxon rank sum test was used to compare all continuous variables and the Mann-Whitney test was used to compare categorical variables.

|  | TB-IRIS (n=25) | Non-IRIS (n=18) | p-value |
| --- | --- | --- | --- |
| Age [Median (IQR)] (years) | 34 (22-52) | 33 (24-55) | ns |
| Female sex [n (%)] | 13 (52%) | 12 (66%) |  |
| Previous TB [n (%)] | 15 (60%) | 10 (55%) |  |
| TB type [n (%)] |  |  |  |
| PTB | 4 (16%) | 2 (11%) |  |
| EPTB | 4 (16%) | 5 (27%) |  |
| EPTB and PTB | 17 (68%) | 11 (61%) |  |
| TB meningitis/neuroTB [n (%)] | 7 (23%) | 4 (21%) |  |
| TB confirmation [n (%)] |  |  |  |
| Cultured Mtb | 9 (36%) | 6 (33%) |  |
| Smear | 6 (24%) | 2 (11%) |  |
| Clinicoradiological | 10 (40%) | 10 (55%) |  |
| Hb [Median (IQR)] (g/dL) | 9.1 (6.4-13) | 9.4 (5.9-14.0) | ns |
| CD4 nadir [Median (IQR)] | 49 (11-209) | 70 (4-272) | ns |
| CD4 count (cells/mm<sup>3</sup>) at week 0 [Median (IQR)] | 68 (21-521) | 111 (4-662) | **0.009** |
| CD4 count (cells/mm<sup>3</sup>) at week 4 [Median (IQR)] | 164 (23-556) | 276 (21-514) | ns |
| Log<sub>10</sub> HIV VL at week 0 [Median (IQR)] | 5.73 (3.96-7.78) | 5.8 (4.21-7.15) | ns |
| Log<sub>10</sub> HIV VL at week 4 [Median (IQR)] | 2.72 (0-3.88) | 2.68 (1.32-3.3) | ns |
| Duration TB treatment to ART [Median (IQR)] (days) | 37 (14-99) | 32 (13-91) | ns |
| Duration ART to TB-IRIS [Median (IQR)] (days) | 15 (4-49) |  |  |
| On steroid treatment at week 0 [n (%)] | 7 (28) | 4 (22) | ns |

### Expansion of Mtb-specific CD4+ T cells at TB-IRIS onset

For phenotypic analyses, we first compared the magnitude of Mtb-specific IFNγ+CD4+ T cell responses between the patient groups before the initiation of ART (Baseline), at week 2, 4 and 6 on ART. Representative examples of IFNγ production by CD4+ T cells following Mtb300 stimulation are presented in **Figure 2A**. We observed no differences in the frequency of Mtb-specific IFNγ+CD4+ T cells between the two groups in cross-sectional comparisons at any time point **(Figure 2B)**. However, the fold change in Mtb-specific IFNγ+CD4+ T cell frequency between baseline and week 2 was significantly higher in the TB-IRIS group compared to the non-IRIS group (median fold change: 1.9 [IQR: 0.83-19.3]) and 0.9 [IQR: 0.25-1.6], respectively, p=0.04) **(Figure 2C)**. This significant increase was exclusively observed in TB-IRIS patients between baseline (median: 0.08% [IQR: 0.0-0.2]) and 2 weeks on-ART (median: 0.13% [IQR: 0.0-0.71]) (p=0.039) **(Supplementary Figure 2)**. We next investigated the phenotype of Mtb-specific IFNγ+CD4+ T cell that could potentially characterise the role of these cells in the pathogenesis of TB-IRIS in humans.

**Figure 2.**
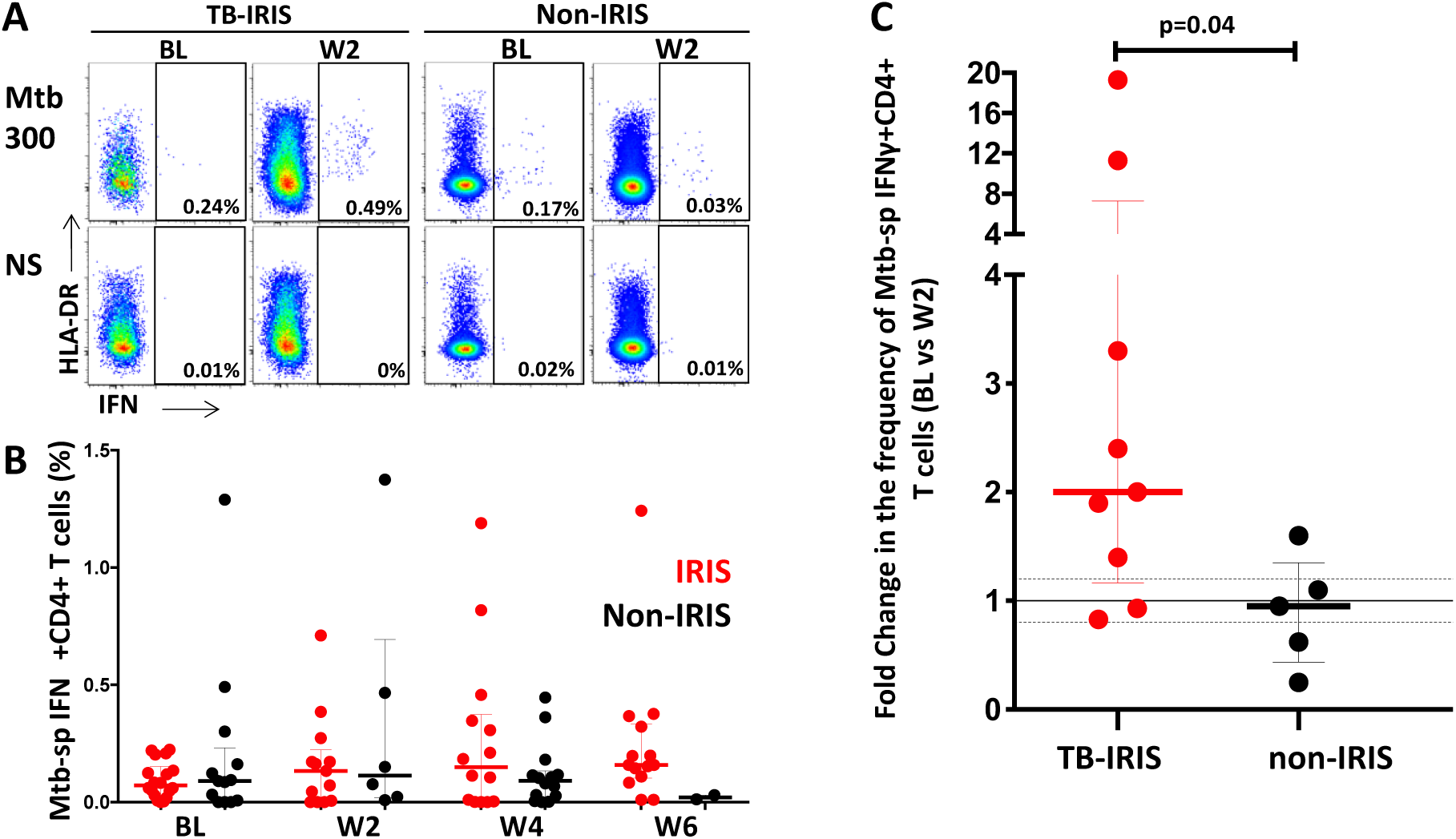
Frequencies of Mtb-specific IFNγ+CD4+ T cells in TB-IRIS and non-IRIS patients. **A**, Representative flow plots of IFNγ production in response to Mtb peptide pool (Mtb300) and non-stimulated controls (NS) at baseline (BL, prior to initiation of antiretroviral therapy, ART) and 2 weeks on ART (W2). **B**, Frequencies of IFNγ producing CD4+ T cells in TB-IRIS (red) from baseline (BL, n= 16), through 2 weeks (W2, n= 9), 4 weeks (W4, n= 10) and 6 weeks (W6, n=12) and non-IRIS (black) from BL= 11, through W2, n= 4, W4, n= 8 and W6, n= 1 on ART. **C**, Fold change in the frequency of IFNγ+CD4+ T cells in TB-IRIS and non-IRIS patients between baseline (prior to ART) and 2 weeks on ART. The Wilcoxon ranked test was used for the statistical comparison of paired samples and the Mann-Whitney-U test was used for unpaired samples. Only statistically significant data with a p value of 0.05 or less are indicated on graphs

### No differences in the expression of Eomes or Tbet between patients with and without TB-IRIS at any tested time point

Based on our mouse model data and a recent report by Hsu *et al*. reporting that Eomes was significantly upregulated over Tbet in MAC-specific IFNγ+CD4 T cells of MAC-IRIS patients at disease onset [26], we determined whether these transcription factors were differentially expressed between TB-IRIS and non-IRIS patients. We observed no differences in the frequency of Mtb-specific IFNγ+Eomes+CD4+ T cells between the two clinical groups at baseline or any time point on ART (**Supplementary Figure 3**).

Eomes and Tbet expression in Mtb-specific IFNγ+CD4 T cells were highly variable between patients but not statistically different between the two groups at baseline **(Supplementary Figure 4)** or any other time point (data not shown). The expression of Eomes in Mtb-specific IFNγ+CD4 T cells at baseline was approximately 50% and was comparable between the two groups (**Supplementary Figure 4B)**. Similarly, Tbet expression in Mtb-specific IFNγ+CD4+ T cells was comparable between TB-IRIS and non-IRIS groups; approximately 60% of cells expressed intermediate levels of Tbet (Tbet dim, Tbet+) and 25% expressed high Tbet levels at baseline (Tbet high, Tbet++, **Supplementary Figure 4C&D)**. Furthermore, Eomes expression on total CD4+ T cells in TB-IRIS patients was comparable to non-IRIS controls at all-time points **(Supplementary Figure 6A&B)**. However, we did observe a slight increase in Eomes expression between baseline and week 2 (which corresponds to IRIS onset) in TB-IRIS patients (medians: 4.48% vs 7.6%, respectively, p=0.03). This was not observed in non-IRIS controls **(Supplementary Figure 5C)**. Previous studies have reported a higher frequency of both *M. avium* and Mtb-specific effector memory CD4 T cells in unmasking and paradoxical TB-IRIS patients compared to non-IRIS patients [24, 34]. Furthermore, a positive correlation between CD4+ T cell memory and Eomes expression is well established [27]. Therefore, it is possible that the increase in Eomes expression observed in total CD4 T cells could be related to an expansion of effector cells. Finally, Tbet expression on total CD4+ T cells was comparable between TB-IRIS and non-IRIS patients at baseline with no significant differences observed longitudinally on ART (data not shown). In further analyses, we defined the co-expression of Eomes and Tbet, identifying five Eomes/Tbet subsets: Eomes-Tbet-, Eomes-Tbet+, Eomes+Tbet+, Eomes-Tbet++ and Eomes+Tbet++, as previously described [35] (**Figure 3A**). The distribution of these subpopulations within Mtb-specific IFNγ+CD4+ T cells was comparable between TB-IRIS and non-IRIS groups prior to ART initiation (i.e. baseline) **(Figure 3B)** and longitudinally on ART (data not shown). No significant changes in the distribution of Eomes and Tbet in Mtb-specific IFNγ+CD4+ T cells were observed over time within the two groups (**Figure 3C**).

**Figure 3.**
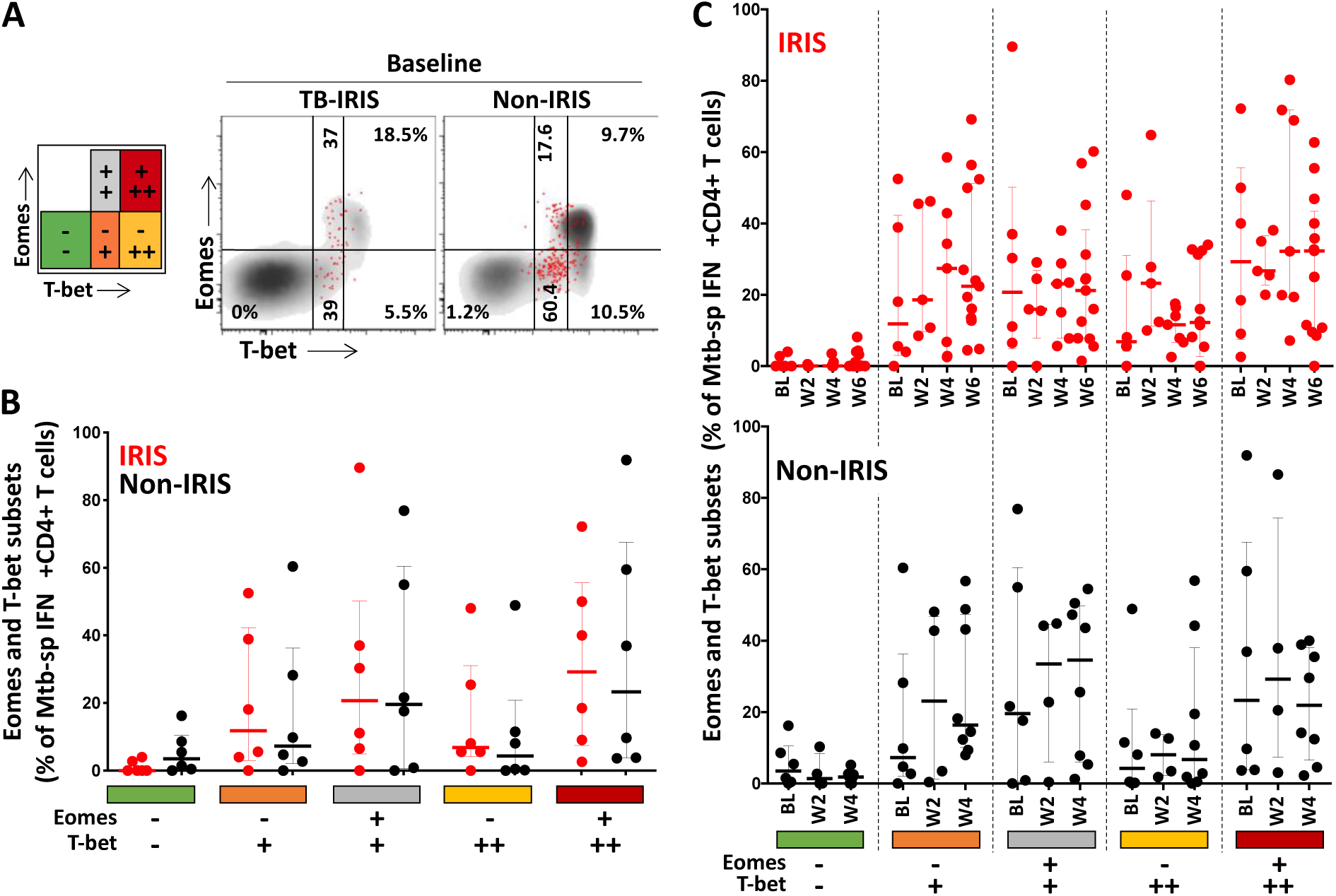
HLA-DR expression on Mtb-specific IFNγ+CD4+ T cells in TB-IRIS and non-IRIS patients. **A**, Representative flow plot of HLA-DR expression on Mtb-specific IFNγ+CD4+ T cells (red) and total CD4+ T cells (black) in one TB-IRIS and one non-IRIS patient at two weeks post ART initiation (W2). **B**, Expression of HLA-DR on Mtb-specific IFNγ+CD4+ T cells in TB-IRIS (red) from baseline (BL, n= 6), through 2 weeks (W2, n= 5), 4 weeks (W4, n= 7) and 6 weeks (W6, n= 13) and non-IRIS patients (black) from baseline (BL, n= 6), through 2 weeks (W2, n= 4), and 4 weeks (W4, n= 8) on-ART. **C**, Frequency of HLA-DR on Mtb-specific IFNγ+CD4+ T cells from baseline to 6 weeks on ART in TB-IRIS and non-IRIS patients. The Wilcoxon ranked test was used for the statistical comparison of paired samples and the Mann-Whitney-U test was used for unpaired samples. Only statistically significant data with a p value of 0.05 or less are indicated on graphs.

However, in total CD4+ T cells there was a significant reduction in Eomes-Tbet-CD4+ T cells between baseline (median: 79.0%, IQR: 21.2-93.1) and week 2 (median: 65.5%, IQR: 15.3-84.4) (p=0.02) and baseline and week 4 (median: 54.5%, IQR: 21.8-94.5) (p=0.009) in TB-IRIS patients. These changes were countered by a progressive and significant increase in the proportion of Eomes-Tbet+ and Eomes+Tbet+ CD4+ T cells over the first 6 weeks of ART in TB-IRIS patients. Conversely, no changes over time were observed in the distribution of any of the Eomes/Tbet subsets in non-IRIS patients (**Supplementary Figure 6C**).

### Elevated HLA-DR expression at the time of TB-IRIS onset compared to non-IRIS controls

To further characterise the phenotype of Mtb-specific IFNγ+CD4+ T cell responses, we compared the activation profile (HLA-DR) and cytotoxic potential (Granzyme B) between TB-IRIS patients and non-IRIS controls. We observed a trend towards high pre-ART HLA-DR expression (p=0.18) in TB-IRIS compared to non-IRIS patients. Responses were characterized by a significantly higher expression of HLA-DR in TB-IRIS compared to non-IRIS patients at 2 weeks on ART (median: 79.3% [IQR: 66-96] and 40.9% [IQR: 27-56], respectively, p=0.016) **(Figure 4A&B)**. No significant changes over time were observed when data were analysed longitudinally for both groups respectively **(Figure 4C)**.

**Figure 4.**
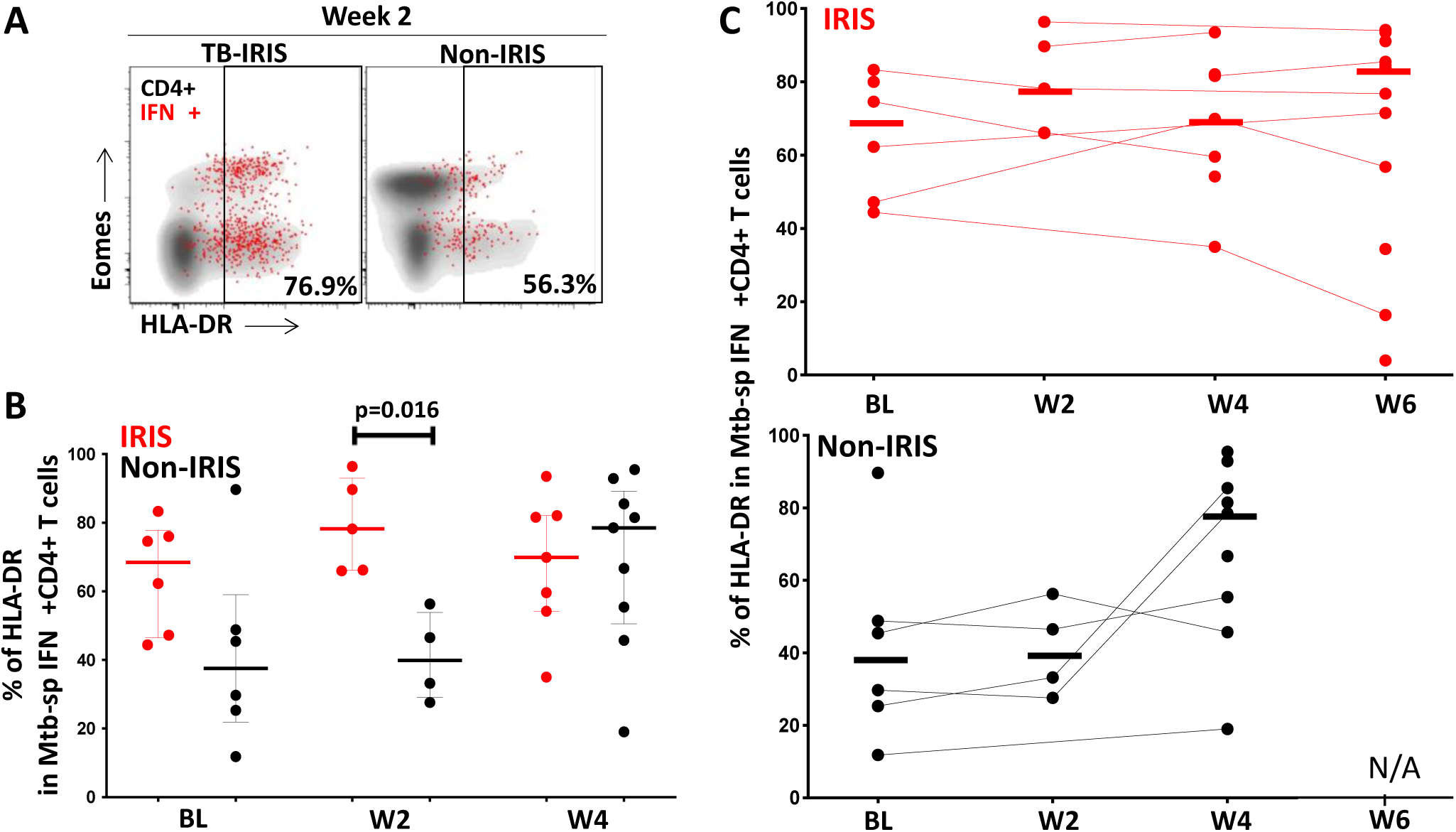
Eomes and T-bet expression profile in Mtb-specific IFNγ+CD4+ T cells in TB-IRIS and non-IRIS patients. **A**, Representative flow plot of Eomes and T-bet expression on Mtb-specific IFNγ+CD4+ T cells (red) and total CD4+ T cells (black) in one TB-IRIS and one non-IRIS patient prior to ART initiation (BL). **B**, Distribution of Mtb-specific IFNγ+CD4+ T cells amongst distinct Eomes and T-bet subsets: (Eomes-T-bet-; Eomes-T-bet+; Eomes+ T-bet+; Eomes-T-bet++; Eomes+ T-bet++) in TB-IRIS (red, n= 6) and non-IRIS patients (black, n= 6) at BL. **C**, Evolution of Eomes and T-bet profile in Mtb-specific IFNγ+CD4+ T cells in TB-IRIS from BL, (n= 6), through 2 weeks (W2, n= 5), 4 weeks (W4, n= 7) and 6 weeks (W6, n= 13) and non-IRIS patients (black) from BL, (n= 6), through 2 weeks (W2, n= 4) and 4 weeks (W4, n= 8) on-ART. The Wilcoxon ranked test was used for the statistical comparison of paired samples and the Mann-Whitney-U test was used for unpaired samples. Only statistically significant data with a p value of 0.05 or less are indicated on graphs.

However, we observed an increase in HLA-DR expression in total CD4+ T cells in TB-IRIS patients between baseline and week 2 (p=0.002) and week 4 (p=0.0005) and non-IRIS patients between baseline and week 4 (p=0.0098) **(Supplementary Figure 7)**.

### Elevated HLA-DR and granzyme B expression in Mtb-specific CD4 T cells co-expressing Eomes and Tbet in patients with TB-IRIS compared to non-IRIS controls

Finally, we investigated the activation and cytotoxic potential of Mtb-specific IFNγ+CD4+ T cells in relation to their transcription factor profile at the time of IRIS onset (Week 2). While HLA-DR expression was comparable across the different Eomes and Tbet subsets in both groups, HLA-DR expression was higher in TB-IRIS compared with non-IRIS patients in specific Eomes/Tbet subsets, including Eomes+Tbet+ (median: 83.9% vs 57.9%, respectively; p=0.032) and Eomes-Tbet++ (median: 83.3% vs 36.4%, respectively; p=0.032) (**Figure 5A**).

**Figure 5.**
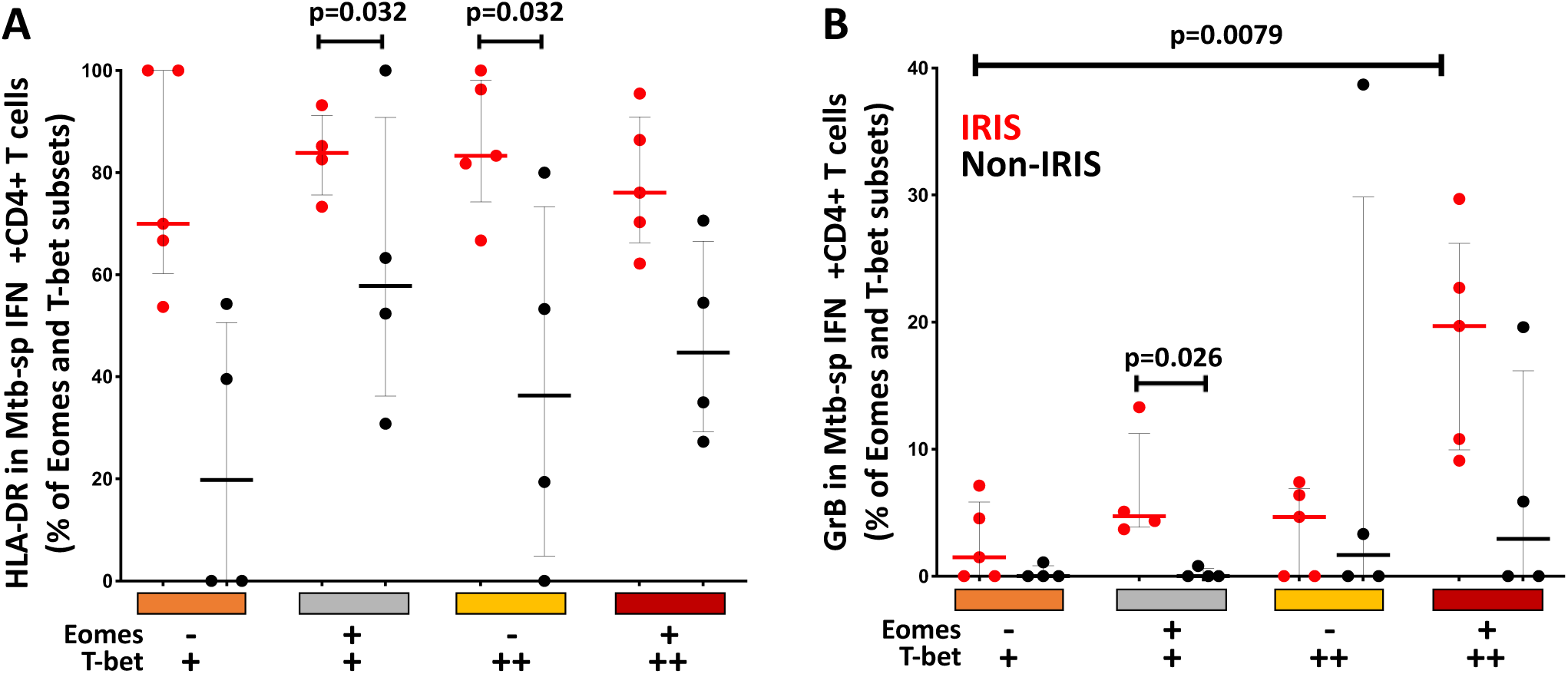
Expression of HLA-DR and granzyme B on Eomes and T-bet expressing subsets of Mtb-specific IFNγ+CD4+ T cells two weeks on ART. **A**, Expression of HLA-DR and **B**, granzyme B on Eomes and T-bet subsets (Eomes-, T-bet+, Eomes+, T-bet+, Eomes-, T-bet++ and Eomes+, T-bet++) of Mtb-specific IFNγ+CD4+ T cells in TB-IRIS (red, n= 5), and non-IRIS patients (black, n= 4), 2 weeks on ART. The Mann-Whitney-U test was used for statistical comparison of unpaired samples. Only statistically significant data with a p value of 0.05 or less are indicated on graphs.

Notably, no differences in Granzyme B expression were observed in IFNγ+CD4 T cells between the two groups in cross-sectional comparisons **(Supplementary Figure 8)**. However, Granzyme B expression was significantly higher in Eomes+Tbet+ Mtb-specific IFNγ+CD4+ T cells in patients with TB-IRIS compared to non-IRIS controls at week 2 on ART (median: 4.7% vs 0%, respectively; p=0.026). There was also a trend towards higher Granzyme B expression in the Eomes+Tbet++ Mtb-specific IFNγ+CD4+ subset in patients with TB-IRIS compared to non-IRIS controls at week 2 (median: 19.7% vs 2.9%, respectively; p=0.063) **(Figure 5B)**. This trend was not observed at other time points (data not shown).

## DISCUSSION

Hsu et al. recently reported that in HIV-1 and *M. avium* co-infected patients, *M. avium*-specific IFNγ+CD4+ T cells were characterized by higher expression of Eomes than Tbet at IRIS onset [26], suggesting potential involvement of Eomes in mycobacterial IRIS pathogenesis. While the functional role of Eomes is well established in CD8 T cells [27, 28], its role in CD4 T cells is less clear. Some reports implicate its expression in the pathogenesis of chronic inflammatory disorders [36–38], while others suggest a regulatory role in T cells [39]. Therefore, to define whether aberrant expression of transcription factors in CD4 T cell associate with the development of IRIS, we investigated the role of Eomes and Tbet in a experimentally-induced MAC-IRIS mouse model and compared the phenotype of Mtb-specific IFNγ+CD4+ T cells between HIV-associated TB patients who developed TB-IRIS and those who did not.

The MAC-IRIS mouse model showed that mimicking T cell reconstitution using Eomes knock-out CD4 T cells led to enhanced mice survival compared to wildtype, supporting the hypothesis that Eomes expression in CD4 T cells could play a role in IRIS pathogenesis [26]. However, while we demonstrated that Mtb-specific IFNγ+CD4+ T cells from TB-IRIS patients expressed high Eomes levels (~ 50%) that are comparable to those reported by Hsu et al. [26], we did not observe any difference in Eomes expression between TB-IRIS and non-IRIS patients. This suggests that the expression of Eomes expression in Mtb-specific CD4 T cells (or overall CD4 compartment) on its own does not predict nor characterize TB-IRIS pathogenesis.

The expression profile of Tbet in CD4+ T cells in this cohort mirrors that described by Knox *et al*. [40], where three distinct populations were discernible. Most Mtb-specific IFNγ+CD4+ T cells expressed Tbet with ~ 65% being Tbet dim and ~ 25% Tbet bright. Moreover, we found no significant differences in the co-expression profile of Eomes and Tbet in Mtb-specific IFNγ+CD4+ T cells at TB-IRIS onset or at other time points between the two clinical groups. However, the distribution of Eomes and Tbet subsets in total CD4+ T cells were altered during the course of ART with increasing expression of both Tbet+ and Eomes/Tbet co-expressing CD4+ T cells in TB-IRIS patients on ART. Further studies are needed to confirm these observations and define their relevance.

In this cohort, TB-IRIS patients had significantly lower blood CD4 T cell counts compared to non-IRIS patients at baseline, as previously described [8, 41] and we observed a significant expansion in the frequency of Mtb-specific IFNγ+CD4+ T cells 2 weeks after the initiation of ART. Recently, Vignesh *et al*. described elevated pre-ART frequencies of Mtb-specific CD4 T cell responses which further expanded in TB-IRIS patients at disease onset [41]. We did not observe such differences at baseline in this or previous studies [23]. Clinical differences between the cohorts might account for these discrepancies.

Several studies have demonstrated that TB-IRIS is characterized by an increase in mycobacteria-specific CD4 T cell responses at disease onset [22, 41–44]. However, increased mycobacterial-specific CD4 T cell frequencies following ART is not systematically observed in all TB-IRIS patients, and pathogen-specific CD4 T cell expansion can also be observed in some non-IRIS patients [23]. This suggests that Mtb-specific CD4 T cell reconstitution upon ART is not be the only mechanism involved in TB-IRIS.

To further elucidate the contribution of Mtb-specific IFNγ+CD4+ T cells in TB-IRIS pathology, we characterised their phenotype in TB-IRIS patients. We demonstrated that Mtb-specific IFNγ+CD4 T cells of TB-IRIS patients had elevated HLA-DR expression prior to the initiation of ART and this was significantly upregulated in TB-IRIS patients at week 2 on ART compared to non-IRIS patients. Similarly, others have demonstrated that Mtb-specific CD4 T cells are activated [24], and polyfunctional [25, 42], compared to non-IRIS controls at IRIS onset.

Consistent with our previous findings [23], we did not observe any significant differences in the expression of HLA-DR in total CD4+ T cells between the two clinical groups over time in a cross section analysis. However, like Antonelli *et al*, we observed increased HLA-DR expression in total CD4+ T cells of TB-IRIS patients from baseline to week 2 and 4 [24]. Similar observations were reported by Haridas et al. at the time of IRIS onset [45].

Lastly, Granzyme B expression was enriched in Eomes/Tbet co-expressing Mtb-specific IFNγ+CD4+ T cells at 2 weeks on ART in TB-IRIS patients. Although this represents a modest proportion of Eomes+Tbet+ cells, this is consistent with mouse data from experimental autoimmune encephalitis showing the capacity of Eomes+ IFNγ+CD4 T cells to acquire cytotoxic attributes [36]. Moreover, our group has previously shown TB-IRIS to be associated with increased transcript abundance and secretion of granzyme B by PBMC of TB-IRIS patients at week 2 on ART [46].

There were several limitations to this study. The number of samples analysed were limited, consequently, larger cohort studies are needed to verify these findings. We assessed responses in peripheral blood when clinical manifestations are often localized in tissues. Finally, several patients with severe disease received corticosteroids prior to or while on ART. Our previous findings however, suggest that corticosteroid treatment does not have a significant impact on *ex vivo* T cell functional responses in TB-IRIS patients [47].

In conclusion, while the mouse model data suggested that CD4 T cell expression of Eomes promotes IRIS, there were no differences in the expression of Eomes or Tbet transcription factor in Mtb-specific IFNγ+ CD4 T cells between patients who developed TB-IRIS and non-IRIS controls. We found that TB-IRIS was associated with an increase of Mtb-specific CD4 T cells at onset. Moreover, increased expression of markers of immune activation and cytotoxicity in Mtb-specific CD4 T cell subsets in TB-IRIS patients suggests these cells may contribute to pathogenesis of TB-IRIS. Improved understanding of the pathophysiology of IRIS should enable the development of new diagnostic tools and better targeted treatments.

## Acknowledgements

The authors thank the study participants, the clinical staff and the laboratory staff involved in this study.

---

**Supplementary Table 1.** Individual patient duration to first TB-IRIS episode, symptom manifestation and steroid management. Different patients were on different combination of ART regimen including Stavudine (D4T), Lamivudine (3TC), Efavirenz (EFZ), Zidovudine (AZT) and Tenofovir (TDF).

| IRIS patient Identifier | ART regimen | Duration ART Start to TB-IRIS (days) | TB-IRIS system involvement | Steroids to Treat TB-IRIS | Duration from TB-IRIS onset to starting steroids (days) |
| --- | --- | --- | --- | --- | --- |
| IR 4011-3 | D4T, 3TC, EFZ | 6 | Pulmonary, nodal, effusion and abdominal | Yes | 9 |
| IR 4035-1 | AZT, 3TC, EFZ | 42 | Nodal, abdominal | No |  |
| IR 4074-8 | TDF, 3TC, EFZ | 10 | Pulmonary, nodal | Yes | 5 |
| IR 4071-5 | TDF, 3TC, EFZ | 8 | Pulmonary, effusion, abdominal | No |  |
| IR 4075-9 | D4T, 3TC, EFZ | 12 | Abdominal | Yes | na |
| IR 4076-0 | TDF, 3TC, EFZ | 9 | Nodal | No |  |
| IR 4081-7 | TDF, 3TC, EFZ | 10 | Abdominal | No |  |
| IR 4108-1 | TDF, 3TC, EFZ | 9 | Pulmonary, abdominal | No |  |
| IR 4018-0 | D4T, 3TC, EFZ | 13 | Pulmonary | Yes | 4 |
| IR 4020-4 | D4T, 3TC, EFZ | 30 | Nodal | Yes | 13 |
| IR 4047-5 | D4T, 3TC, EFZ | 8 | Pulmonary, abdominal | Yes | 5 |
| IR 4110-5 | D4T, 3TC, EFZ | 4 | Pulmonary, abdominal | No |  |
| IR 4052-2 | TDF, 3TC, EFZ | 13 | Pulmonary, abdominal | No |  |
| IR 4078-2 | TDF, 3TC, EFZ | 26 | Neurological, pulmonary, abdominal | Yes | 20 |
| IR 4096-4 | TDF, 3TC, EFZ | 14 | Pulmonary, abdominal | No |  |
| IR 4115-0 | AZT, 3TC, EFZ | 26 | Pulmonary | Yes | 16 |
| IR 4010-2 | D4T, 3TC, EFZ | 7 | Nodal, abdominal | Yes | 56 |
| IR 4015-7 | D4T, 3TC, EFZ | 15 | Pulmonary, abdominal | Yes | 20 |
| IR 4021-8 | D4T, 3TC, EFZ | 5 | Pulmonary | Yes | 2 |
| IR 4036-2 | D4T, 3TC, EFZ | 7 | Nodal, neurological | Yes | 1 |
| IR 4080-6 | TDF, 3TC, EFZ | 11 | Neurological | Yes | na |
| IR 4085-1 | TDF, 3TC, EFZ | 14 | Articular | Yes | na |
| IR 4088-4 | TDF, 3TC, EFZ | 49 | Neurological | Yes | 9 |
| IR 4095-3 | TDF, 3TC, EFZ | 12 | Pulmonary, abdominal | Yes | 3 |
| IR 4111-6 | TDF, 3TC, EFZ | 8 | Abdominal | Yes | 8 |

## FIGURE LEGENDS

**Supplementary Figure 1.**
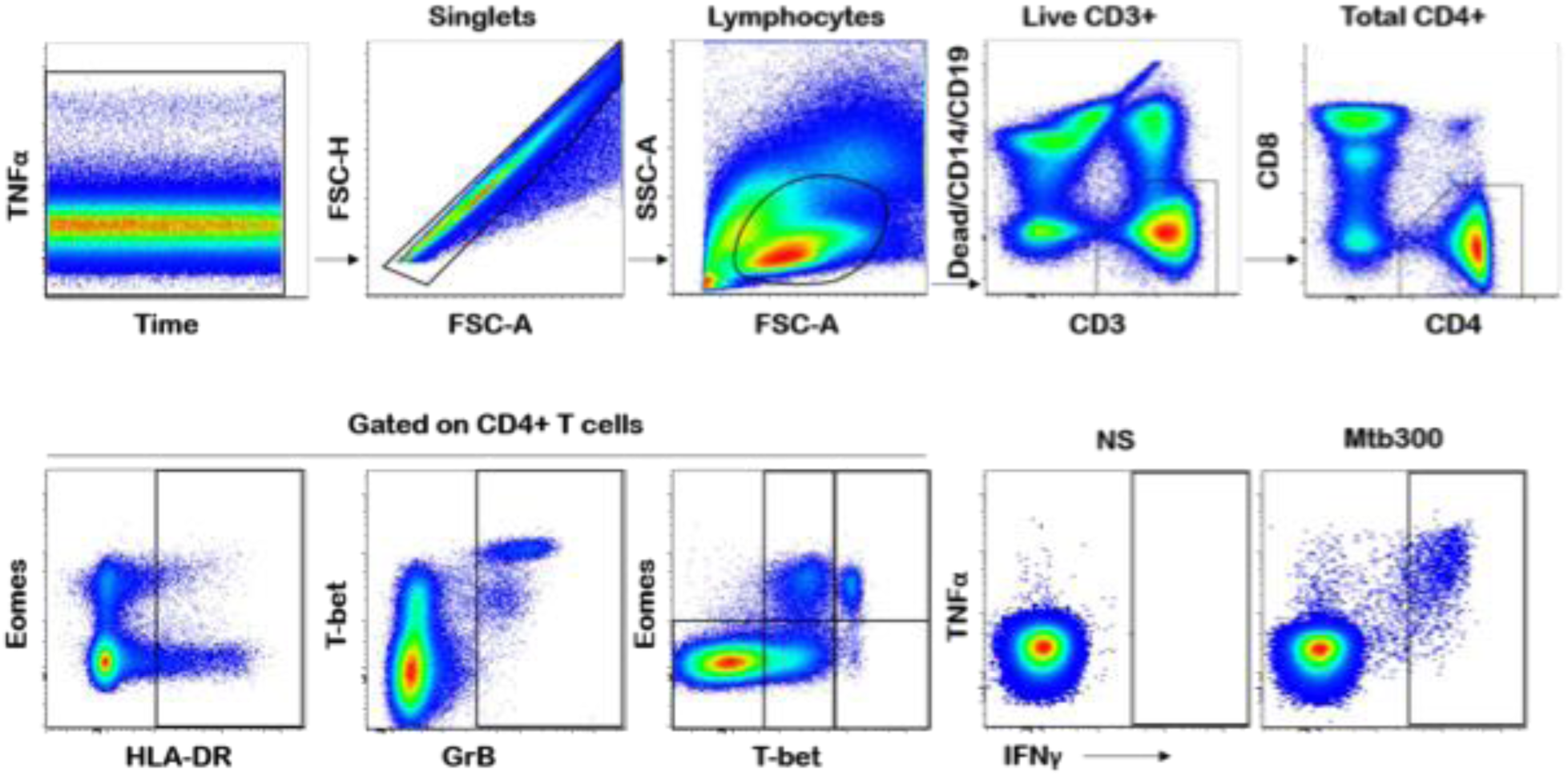
Representative gating strategy for the phenotypic characterization of IFNγ+CD4+ T cells. To phenotype Mtb-specific IFNγ+CD4+ T cell responses, we gated on singlets (FSC-H vs FSC-A), lymphocytes (SSC-A vs FSC-A), live CD3+ cells (dead cells vs live CD3+) and on total CD4+ T cells.

**Supplementary Figure 2.**
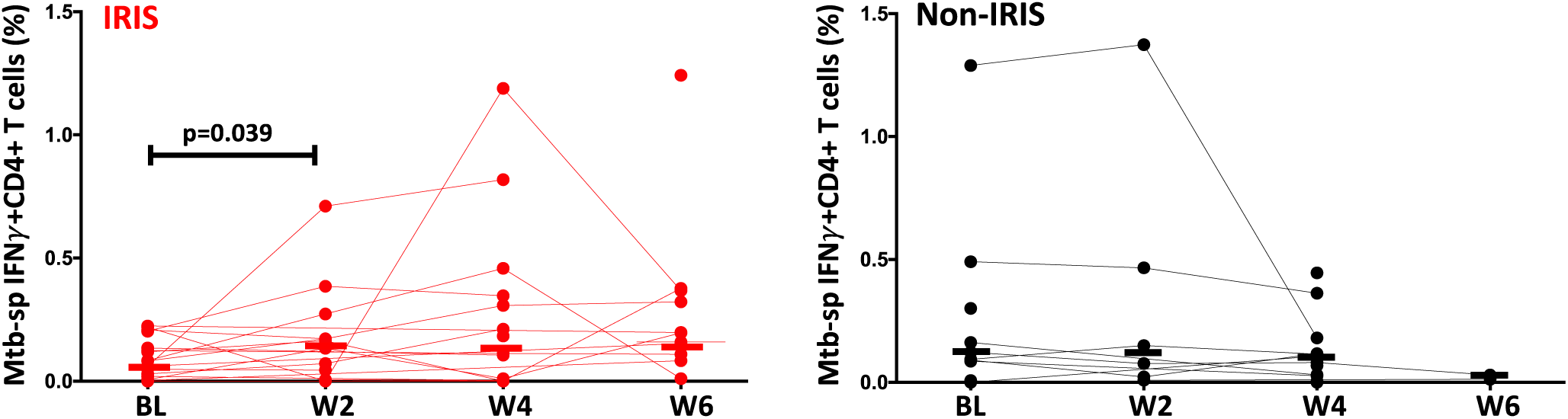
Frequencies of IFNγ producing CD4+ T cells in response to Mtb peptide pool (Mtb300) stimulation in TB-IRIS (red) at baseline (BL, n= 16), 2 weeks (W2, n= 9), 4 weeks (W4, n= 10) and 6 weeks (W6, n=12) and non-IRIS (black) at BL= 11, W2, n= 4, W4, n= 8 and W6, n= 1 after ART initiation. The Wilcoxon ranked test was used for all statistical comparisons. Only statistically significant data with a p value of 0.05 or less are indicated on graphs.

**Supplementary Figure 3.**
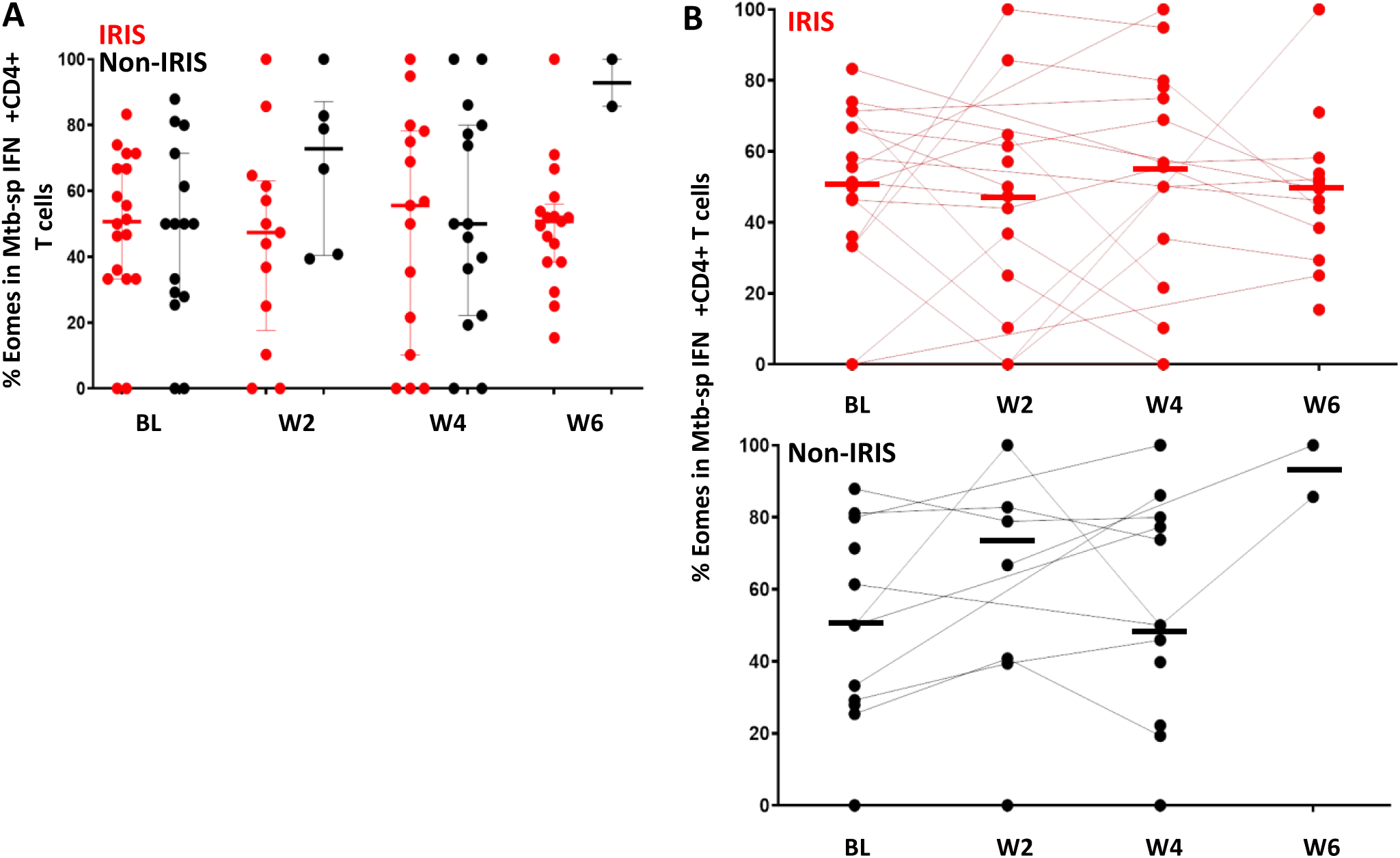
Frequencies of Mtb-specific Eomes+IFNγ+CD4+ T cells in TB-IRIS and non-IRIS patients in response to Mtb peptide pool (Mtb300) stimulation. **A**, Frequencies of Mtb-specific Eomes+ IFNγ+CD4+ T cells in patients with and without TB-IRIS (red and black circles respectively) at baseline (BL, n=18, and 13), 2 weeks (W2, n=13, and 6), 4 weeks (W4, n=14 and 13) and 6 weeks (W6, n=13 and 2) on ART. **B**, Frequencies of IFNγ producing Eomes+CD4+ T cells in TB-IRIS (red) at baseline (BL, n= 18), 2 weeks (W2, n= 13), 4 weeks (W4, n= 14) and 6 weeks (W6, n=13) and non-IRIS (black) at BL= 13, W2, n= 6, W4, n= 13 and W6, n= 2 on ART. The Wilcoxon ranked test was used for all statistical comparisons. Only statistically significant data with a p value of 0.05 or less are indicated on graphs.

**Supplementary Figure 4.**
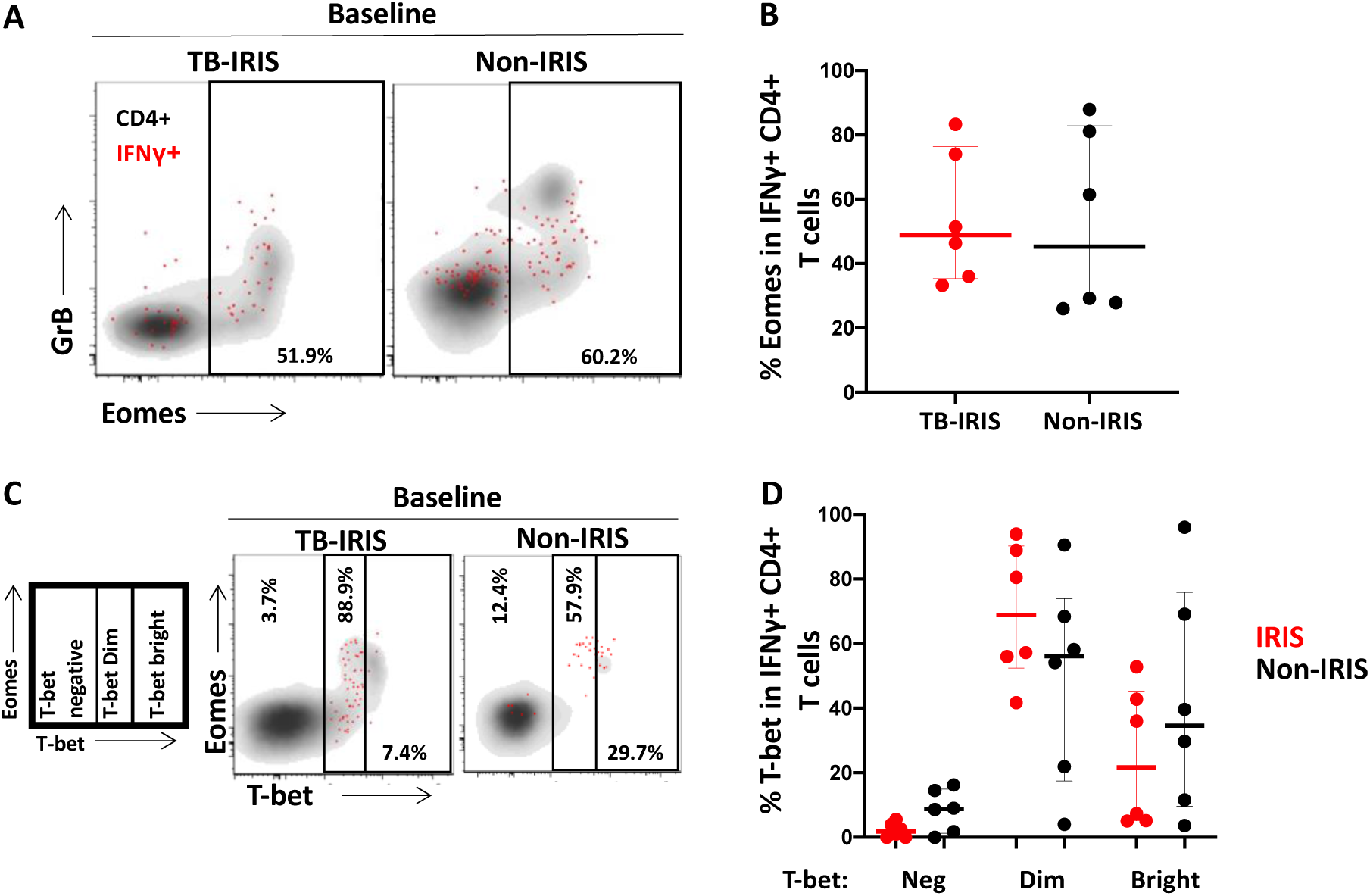
Eomes and T-bet expression on Mtb-specific IFNγ+CD4+ T cells in patients with and without TB-IRIS prior to initiation of antiretroviral therapy (ART) (BL). A, Representative flow plot of the expression of Eomes on Mtb-specific IFNγ+CD4+ T cells (red) and total CD4+ T cells (black) in one TB-IRIS and one non-IRIS patient at BL. B, Summary plot of Eomes expression in Mtb-specific IFNγ+CD4+ T cells between TB-IRIS (n= 6) and non-IRIS patients (n= 6) at BL. C, Representative flow plot of the expression of differentiated T-bet subpopulations on Mtb-specific IFNγ+CD4+ T cells (red) and total CD4+ T cells (black) in one TB-IRIS and one non-IRIS patient at baseline. D, Summary plot of the T-bet expression in Mtb-specific IFNγ+CD4+ T cells between TB-IRIS (n= 6) and non-IRIS patients (n= 6) at BL. The Wilcoxon ranked test was used for all statistical comparisons. Only statistically significant data with a p value of 0.05 or less are indicated on graphs.

**Supplementary Figure 5.**
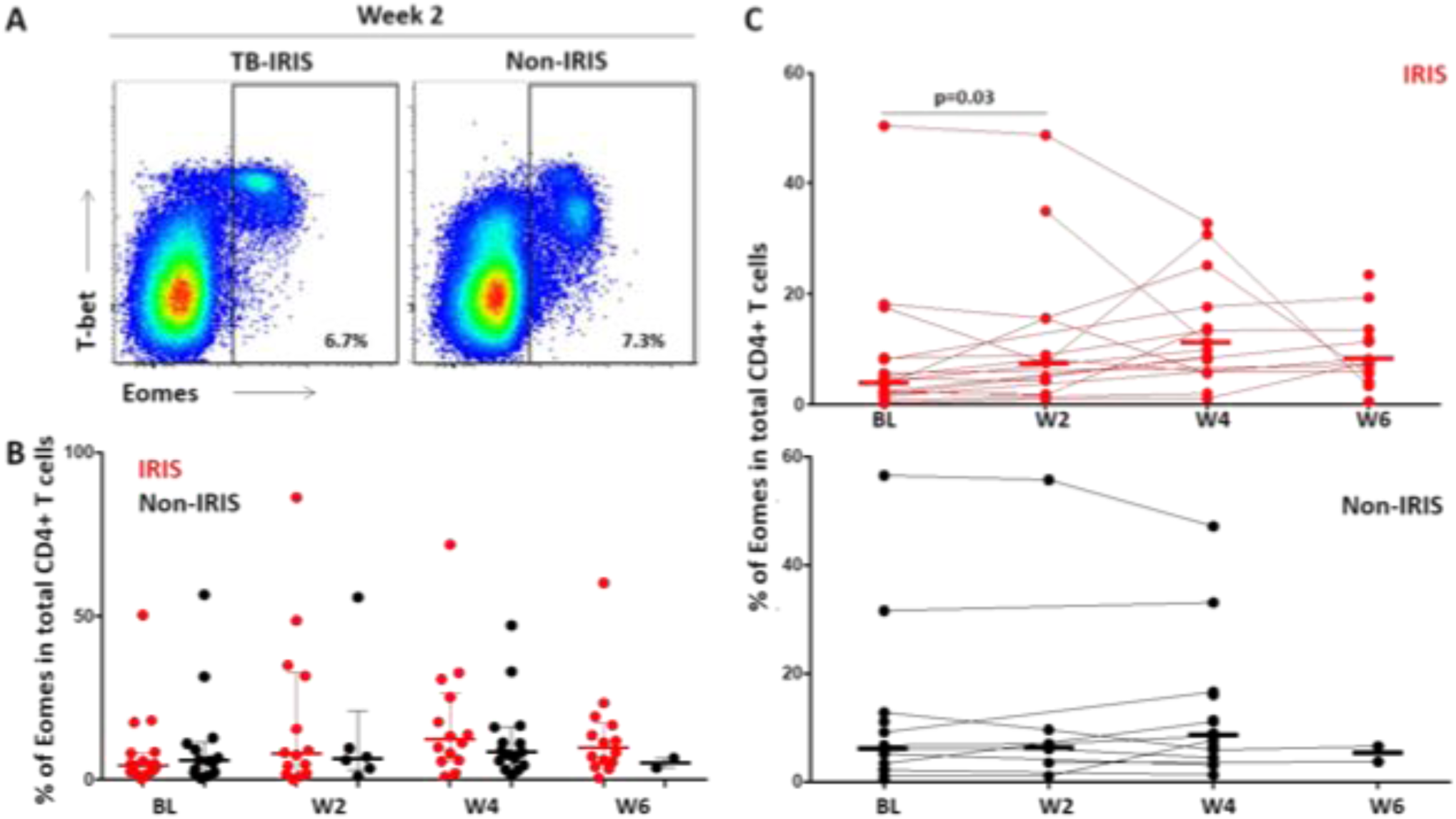
Eomes expression in total CD4+ T cells in patients with and without TB-IRIS. A, Representative flow plot of Eomes expression in one patient with TB-IRIS and one non-IRIS patient in total CD4+ T cells two weeks post ART initiation. B, Cross sectional analyses of Eomes expression in total CD4+ T cells in patients with and without TB-IRIS at baseline (BL, n=18, and 13, respectively), 2 weeks (W2, n=13, and 6), 4 weeks (W4, n=14 and 13) and 6 weeks (W6, n=13 and 2) post-ART. C, Longitudinal analyses of the expression of Eomes in total CD4+ T cells in patients with TB-IRIS (top panel) and non-IRIS controls (bottom panel) from BL to 6 weeks post ART. The Wilcoxon ranked test was used for all statistical comparisons. Only statistically significant data with a p value of 0.05 or less are indicated on graphs.

**Supplementary Figure 6.**
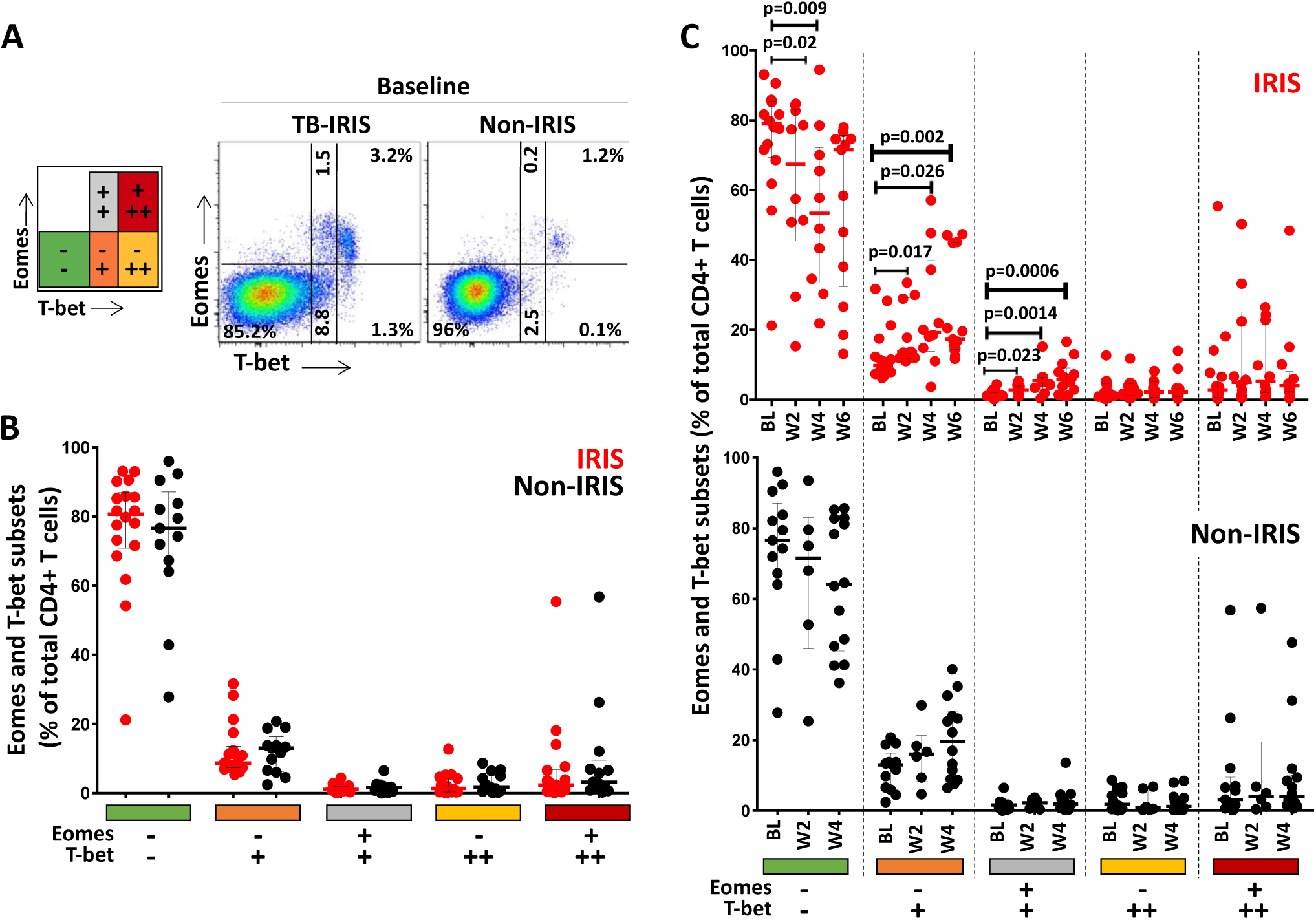
Eomes and T-bet co-expression in total CD4+ T cells in patients with and without TB-IRIS. A, Representative flow plot of Eomes and T-bet co-expression in one patient with TB-IRIS and one non-IRIS patient in total CD4+ T cells two weeks on ART initiation. B, Cross sectional analyses of Eomes and T-bet co-expression in total CD4+ T cells in patients with and without TB-IRIS at baseline (BL, n=18, and 13, respectively), 2 weeks (W2, n=13, and 6), 4 weeks (W4, n=14 and 13) and 6 weeks (W6, n=13 and 2) on ART. C, Longitudinal analyses of the co-expression of Eomes and T-bet in total CD4+ T cells in patients with TB-IRIS (top panel) and non-IRIS controls (bottom panel) from BL to 6 weeks on ART. The Wilcoxon ranked test was used for all statistical comparisons. Only statistically significant data with a p value of 0.05 or less are indicated on graphs.

**Supplementary Figure 7.**
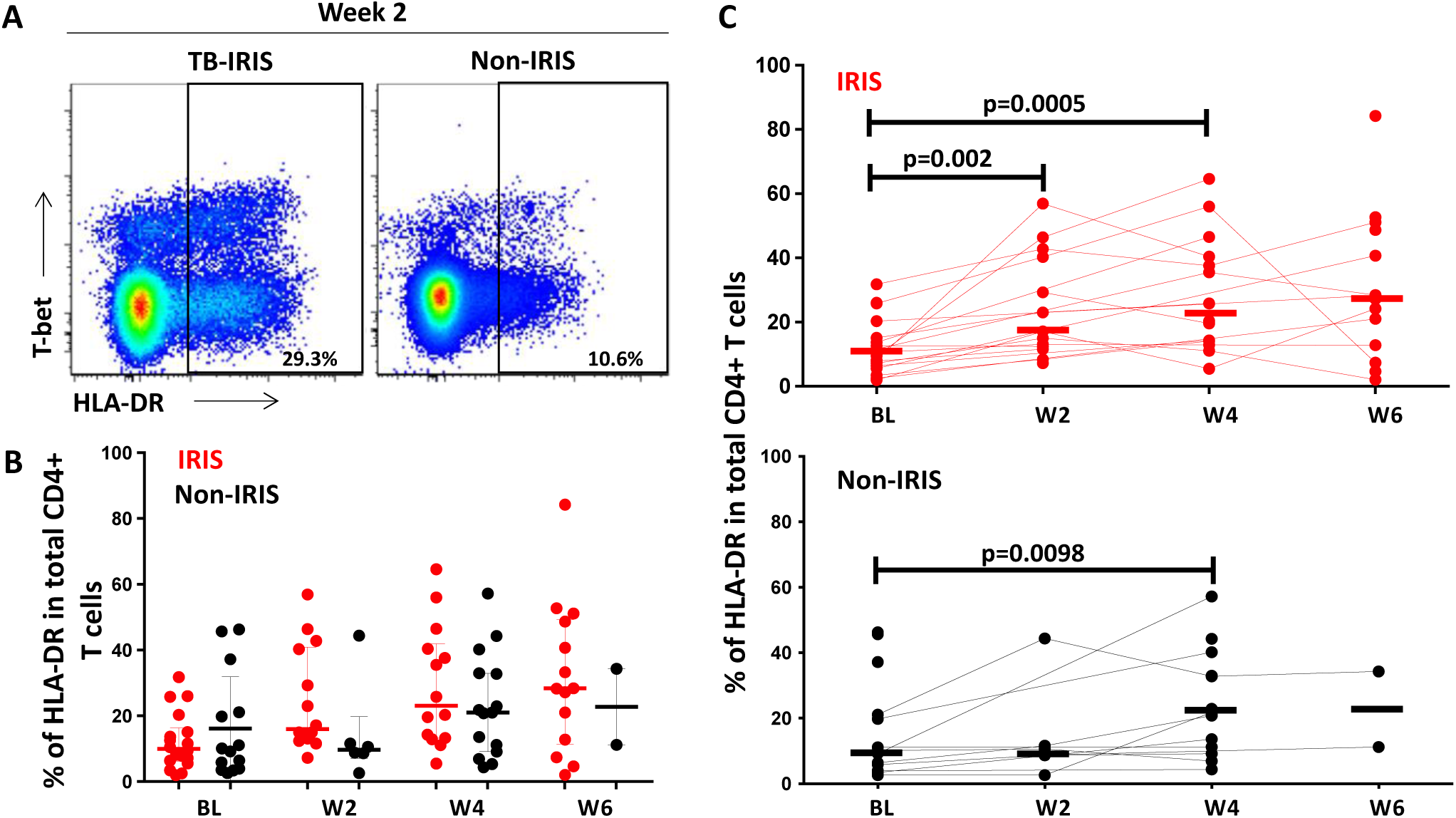
HLA-DR expression in total CD4+ T cells in patients with and without TB-IRIS. A, Representative flow plot of HLA-DR expression in one patient with TB-IRIS and one non-IRIS patient in total CD4+ T cells two weeks post ART initiation. B, Cross sectional analyses of HLA-DR expression in total CD4+ T cells in patients with and without TB-IRIS at baseline (BL, n=18, and 13, respectively), 2 weeks (W2, n=13, and 6), 4 weeks (W4, n=14 and 13) and 6 weeks (W6, n=16 and 2) post-ART. C, Longitudinal analyses of the expression of HLA-DR in total CD4+ T cells in patients with TB-IRIS from BL, n= 18, W2, n= 13,W4, n= 14, W6, n= 13 and non-IRIS controls, BL, n= 13, W2, n= 6, W4, n= 13 and W6, n= 2 post ART. The Wilcoxon ranked test was used for all statistical comparisons. Only statistically significant data with a p value of 0.05 or less are indicated on graphs.

**Supplementary Figure 8.**
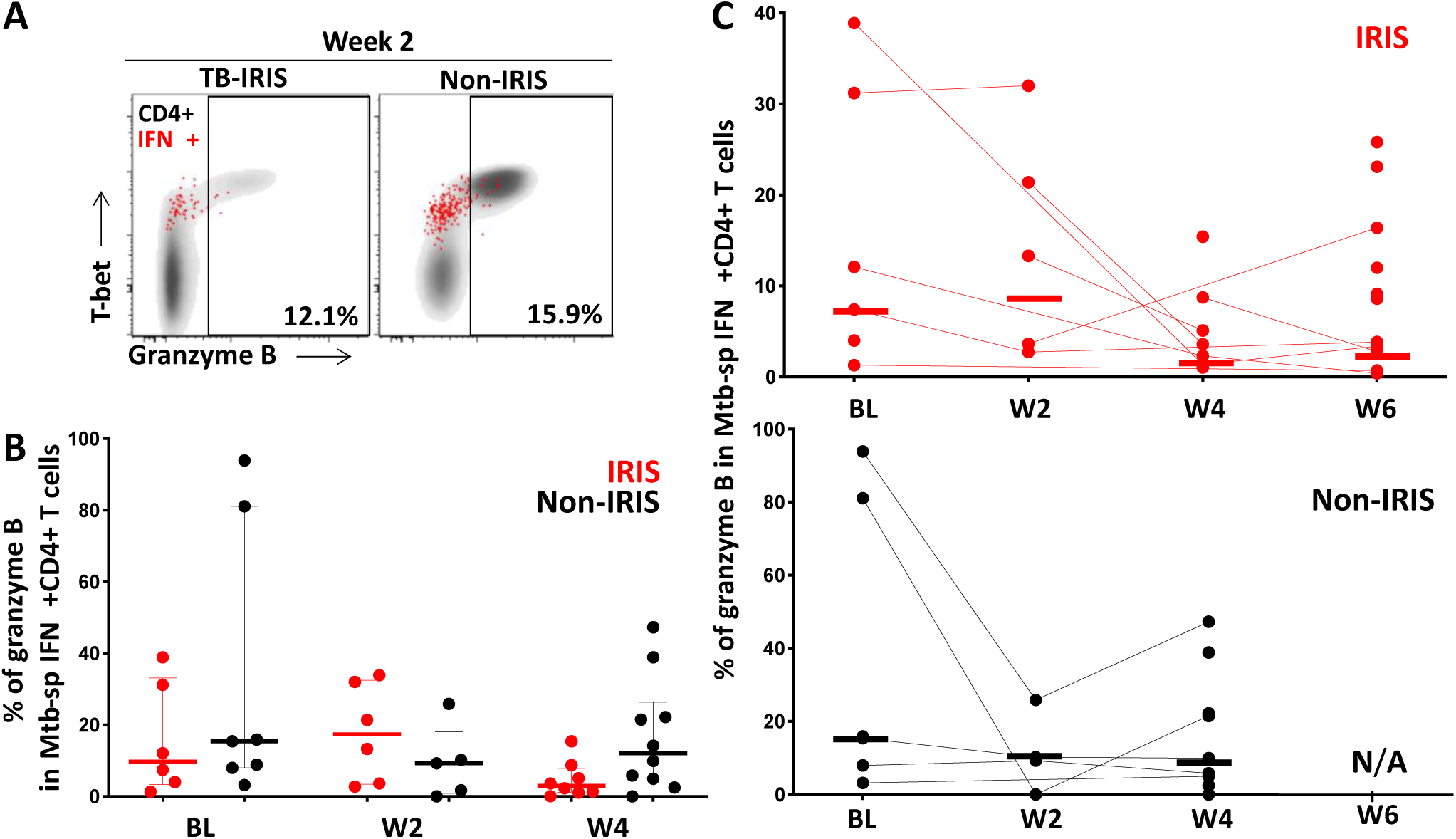
Granzyme B expression in Mtb-specific IFNγ+CD4+ T cells in TB-IRIS and non-IRIS patients. A, Representative flow plot of granzyme B expression in Mtb-specific IFNγ+CD4+ T cells (red) and total CD4+ T cells (gray) in one TB-IRIS and one non-IRIS patient prior to ART initiation (at Baseline, BL). B, Expression of granzyme B in Mtb-specific IFNγ+CD4+ T cells in TB-IRIS (red) at baseline (BL, n= 6), 2 weeks (W2, n= 5), 4 weeks (W4, n= 7) and 6 weeks (W6, n= 13) and non-IRIS patients (black) at baseline (BL, n= 6), 2 weeks (W2, n= 4), and 4 weeks (W4, n= 8) post-ART. C, Expression of granzyme B in Mtb-specific IFNγ+CD4+ T cells from Baseline to 6 weeks post ART in TB-IRIS and non-IRIS patients. The Wilcoxon ranked test was used for all statistical comparisons. Only statistically significant data with a p value of 0.05 or less are indicated on graphs.

